# A mathematical model for cancer dynamics with treatment and saboteur bacteria

**DOI:** 10.1101/2025.04.10.647988

**Authors:** Anna Geretovszky, Gergely Röst

## Abstract

We construct a mathematical model of cancer dynamics with chemotherapeutic treatment, in the presence of bacteria that are capable of metabolizing the chemotherapeutic drug, hence sabotaging the treatment. We investigate the possibility of complementing the cancer treatment with antibiotic drugs, thus eradicating the bacteria or at least mitigating their negative impact on the prospects of therapy. Our model is a nonlinear system of four differential equations, for which we perform a complete analysis, explicitly characterizing the four possible outcomes, depending on whether the cancer cells or the bacteria extinct or persist. Global stability results are proven by the iterative application of a comparison principle, and a bifurcation diagram is created to show the transitions between scenarios with respect to the controllable parameters. We apply our model to an experiment on mice with colon cancer and the drug Gemcitabine.

## 1 Introduction

Cancer remains one of the most pressing challenges in modern medicine. It encompasses a diverse group of diseases characterized by uncontrolled cell division, leading to tumour formation and, in many cases, metastasis to distant organs. Without effective treatment, cancer is fatal, and its global burden continues to rise, partly due to population growth and aging. While advances in medical research, early detection, and risk factor reduction — such as decreased smoking rates — have contributed to a decline in age-standardized cancer mortality rates, current therapies remain ineffective for many patients. In 2025, the global incidence of cancer is projected to reach 21.33 million new cases, with an estimated 10.41 million deaths. Cancer is the second leading cause of mortality worldwide, accounting for approximately 14–19% of all deaths, following cardiovascular diseases ([1, 2, 3]).

Mathematical tools are increasingly being applied in biology and medicine, including oncology ([4, 5, 6]), to aid in the fight against cancer. Over the years, numerous mathematical models have been developed to study cancer dynamics, providing valuable insights into tumour growth, treatment responses, and disease progression. Comprehensive overviews of these models can be found in [7, 8].

There are several strategies for eliminating harmful cells in a patient’s body, one of which is chemotherapy. This treatment employs chemical agents to target the tumour, making it particularly useful for inoperable cases. Chemotherapeutic drugs primarily attack rapidly dividing cells, a hallmark of cancer cells ([9, 10]). However, because these drugs are not selective for cancer cells alone, they also damage healthy tissues, leading to significant side effects. Therefore, optimizing treatment strategies to minimize the overall chemotherapy dosage while maintaining efficacy is crucial. A major challenge in cancer therapy is the development of drug resistance, which can render chemotherapy ineffective. Depending on the type of cancer, there can be different reasons and mechanisms for the emergence of resistance ([9, 11]). One key process is natural selection: cells in the tumour mutate and the ones that acquired hereditary properties that are advantageous against chemotherapy become the majority ([12]). Another mechanism is cellular plasticity, where cells adapt their phenotype in response to environmental cues without genetic mutations. For instance, some cells develop the ability to pump drugs out of themselves using proteins, which is associated with the MDR (multidrug resistant) phenotype ([13]). Additionally, microvesicles, which play a role in intercellular communication between cancer cells, may also contribute to the development of resistance. These microvesicles can transfer resistance-conferring molecules from resistant to non-resistant cells, effectively spreading resistance throughout the tumour, much akin to the spreading of an infection ([14, 15]).

Several mathematical models have been developed to account for drug resistance in cancer treatment ([16]). The motivation for this paper is the findings of the article [17], which describe the relatively recent scientific discovery that microorganisms can influence the effectiveness of chemotherapy treatments targeting cancer tumour cells. Certain (not necessarily malignant) bacteria in the patient’s body can metabolize chemotherapy drugs. Simply speaking, the bacteria “consume” the anti-cancer drug, hence sabotaging the treatment by preventing the drug from reaching the targeted cells, effectively making the tumour resistant to the therapy. In such cases, chemotherapy may need to be supplemented with antibiotics to reduce – or even eliminate – the sabotaging bacterial population, thus opening the way to a more effective chemotherapy treatment. However, excessive antibiotic use must be avoided, as it can further compromise the patient’s already weakened immune system and contribute to antibiotic resistance. Therefore, identifying the optimal combination and dosage of both chemotherapeutic and antibiotic agents is crucial to maximizing treatment efficacy while minimizing adverse effects.

In this paper, we construct a dynamic model to represent the underlying biological mechanisms of tumour-bacteria-drug interactions. The model consists of a nonlinear system of four differential equations that describe the temporal variations of tumour size, the population of drug-metabolizing intra-tumour bacteria, and the concentration of both the chemotherapeutic drug and the antibiotics. To characterize the possible treatment outcomes, we introduce three combined threshold parameters that distinguish four scenarios based on the persistence or eradication of the tumour and the bacterial population. We identify the therapies that achieve the desired outcome, when both the tumour and the bacteria are eradicated. The main result of this paper is that for all four scenarios, we prove the global asymptotic stability of the equilibrium to which the solutions converge. The main tool of the proof is the iterative application of a comparison principle. We also conduct a bifurcation analysis in the parameter space of drug doses, and develop the full bifurcation diagram, demonstrating that single and simultaneous transcritical bifurcations separate these scenarios.

We calibrate a specific version of our model using real-world data, demonstrating its accuracy in capturing key biological dynamics. Finally, we explore potential extensions of our model, including more realistic tumour growth laws and the incorporation of pharmacokinetics and pharmacodynamics. These lead to more complicated systems, but we show that several of our qualitative results remain valid in these extended models.

## 2 Model

### 2.1 Construction

We aim to set up a minimal model that already includes all the relevant biological and pharmacological components. Possible further extensions and more general models will be discussed in Section 4. In constructing our model, we assume a uniform spatial distribution and a well-mixed system, so we use ordinary differential equations. We construct our dynamical model step-by-step, deriving from biological mechanisms, where every variable and parameter has a clear biological interpretation.

Let *T* (*t*) denote the size of the tumour as a function of time. The choice of the growth function is heavily influenced by the specific type of cancer being modeled, as well as the environment in which tumour cells proliferate (e.g., the human body or an in vitro laboratory setting). In [18], various cancer types have been modeled using different functional forms. Among these, the power-law model *T* ^*′*^(*t*) = *rT*^*a*^(*t*) was identified as the most accurate fit in the majority of cases. However, the authors highlight its extreme sensitivity to parameterization, making it unreliable for predictive purposes. Moreover, this model assumes unrestricted tumour growth, contradicting the biological reality that tumour expansion is constrained by factors such as nutrient availability and immune response. Additionally, the parameter *a* lacks direct biological interpretation, meaning the model is not mechanistic. For these reasons, we do not consider this model suitable for our study. In addition to the power-law model, the logistic model performed well, especially for cancer types that are usually treated with the chemotherapeutic drug under investigation in [17]. Hence we use this to describe the variation of the tumour volume:

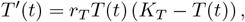

where *r*_*T*_ is the intrinsic tumour growth rate and *K*_*T*_ is the carrying capacity of the environment wrt. the tumour size.

Let *C*(*t*) denote the concentration of the chemotherapy drug as a function of time. In this section, we use a constant drug dose (*γ*) and assume that the drug is absorbed and excreted from the body in proportion to its concentration (with rate *µ*). The appearance of the drug also affects the size of the tumour: assuming mass action law, the cytotoxic effect is proportional to the drug concentration and the number of tumour cells (with rate *δ*). So our system of equations for the tumour and the chemotherapeutic drug is:

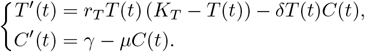

Let *B*(*t*) denote the size of the bacterial population as a function of time. Its growth can also be modelled by a logistic equation, with intrinsic growth rate *r*_*B*_ and carrying capacity *K*_*B*_. The core phenomenon modelled in this paper is that the bacteria can metabolize the chemotherapeutic drug, thus reducing its amount, where again we use mass action law with rate *λ*. The resulting system of equations is

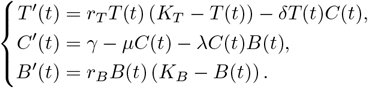

Finally, we introduce the antibiotics that is used to kill the bacteria. Let *A*(*t*) denote the concentration of this as a function of time. Here again, we are working with a constant dose (*α*) and the antibiotic drug is removed with rate *σ*. According to the mass action law, the bacterial population is killed by the antibiotics in proportion to the concentration of the antibiotics and the amount of bacteria (with rate *θ*). This gives our system of equations consisting all essential components:

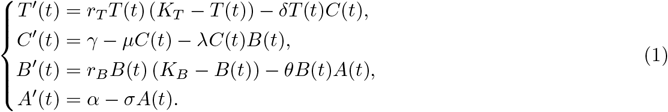

The model variables and the nonnegative parameters are summarized in Table 1.

**Table 1:**
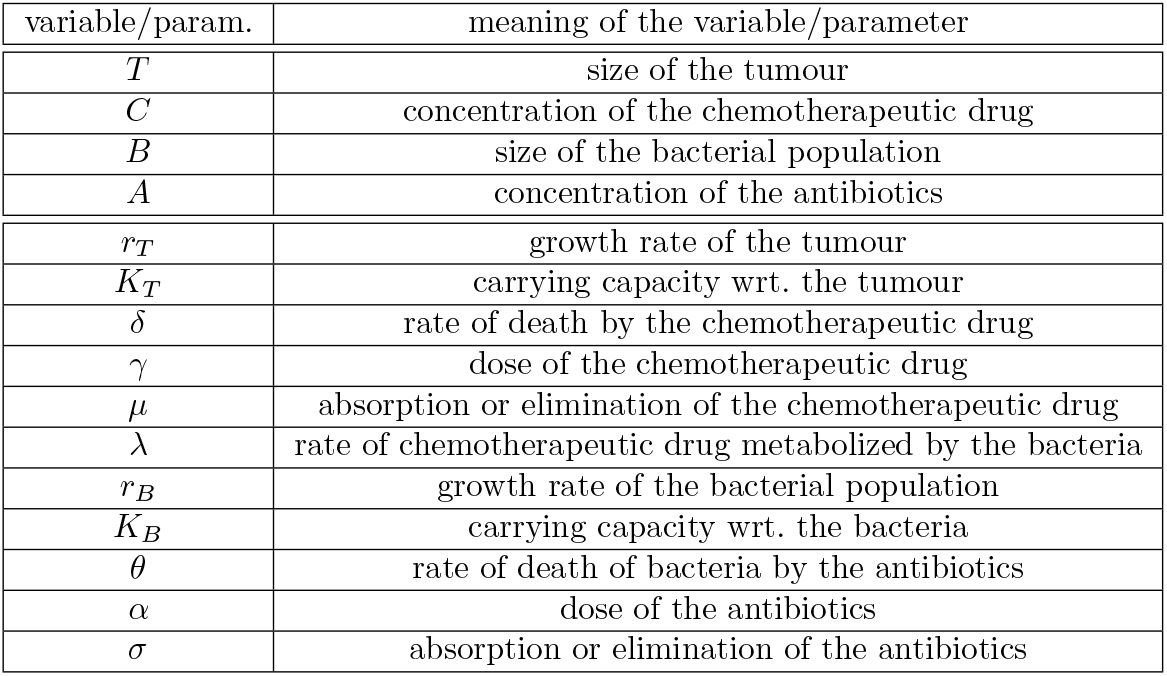
Variables and parameters in the equation system (1).

### 2.2 Analysis

In our model, we have defined variables that describe the size of a population or the concentration of a drug as a function of time. These can obviously only give realistic values if they are non-negative. For this reason, first we show that starting from non-negative initial values, the solutions remain non-negative.

#### Proposition 2.1.

*The equation system* (1) *is well-defined. Starting from nonnegative initial values at t*_0_, *unique solutions exist on the whole* [*t*_0_, *∞*) *interval, moreover, they remain nonnegative and bounded*.

**Proof**. By the Picard-Lindelöf theorem, the equation system (1) has a unique solution. For the nonnegativity part, we use the following well-known statement ([19] Prop. A.1). Let *x*^*′*^(*t*) = *F* (*t, x*) be a differential equation system, where *x*(*t*) = (*x*_1_(*t*), …, *x*_*n*_(*t*)), *F* (*t, x*) = (*F*_1_(*t, x*), …, *F*_*n*_(*t, x*)), and *F* (*t, x*) is defined for all *t ≥*0, ∈ ℝ^*n*^. If the solutions of the initial value problems *x*(*t*_0_) = *x*^0^ are unique for *x*^0^ ∈ [0,*∞*)^*n*^, *t*_0_ *≥*0 and for all *i* = 1, …, *n*

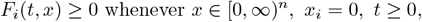

then *x*(*t*) *∈* [0, *∞*)^*n*^ for all *t ≥ t*_0_ *≥* 0, whenever *x*(*t*_0_) *∈* [0, *∞*)^*n*^. *F*_*i*_(*t, x*) *≥* 0 whenever *x ∈* [0, *∞*)^*n*^, *x*_*i*_ = 0, *t ≥* 0, This clearly holds for system (1), hence the solutions remain nonnegative.

The boundedness is concluded from the following observation: when *T* (*t*) *≥ K*_*T*_, then *T* ^*′*^(*t*) *<* 0, which means that lim sup_*t→∞*_ *T* (*t*) *≤ K*_*T*_. The same stands in the case of *C*(*t*), *B*(*T*) and *A*(*t*), with bounds 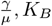and 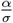, respectively. E.g., 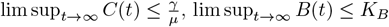 and lim 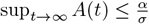.

Since the solutions are bounded, they exist on the whole [*t*_0_, *∞*) interval.

#### Definition 2.2.

Let us define the following compound threshold parameters:

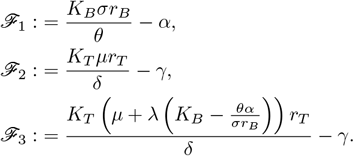

We categorize each equilibrium *E*_*\**_ = (*T*_*\**_, *C*_*\**_, *B*_*\**_, *A*_*\**_) into on of the following types:

- *E*_0_: trivial equilibrium, when both the tumour and the bacteria extinct, i.e. *T*_*\**_ = 0 and *B*_*\**_ = 0;
- *E*_*T*_ : tumour-free equilibrium, when the tumour extincts, but the bacteria persist, i.e. *T*_*\**_ = 0 and *B*_*\**_ *>* 0;
- *E*_*B*_: bacteria-free equilibrium, when the tumour survives, but the bacteria extincts, i.e. *T*_*\**_ *>* 0 and *B*_*\**_ = 0;
- *E*_coex_: coexisting equilibrium, when both the tumour cells and the bacteria persist, *T*_*\**_ *>* 0 and *B*_*\**_ *>* 0.

#### Theorem 2.3

(Equilibrium solutions). *The threshold parameters characterize which types of equilibria of system (1) exist, as follows:*

*Case 1: if ℱ*_1_ *>* 0 *and* ℱ_3_ *>* 0

a. *if* ℱ_2_ *>* 0, *then all four types of equilibria exist;*
b. *if* ℱ_2_ *≤* 0, *then E*_0_, *E*_*T*_ *and E*_*coex*_ *exist;*

*Case 2: if* ℱ_1_ *>* 0 *and* ℱ_3_ *≤* 0, *then E*_0_ *and E*_*T*_ *exist;*

*Case 3: if* ℱ_1_ *≤* 0 *and* ℱ_2_ *>* 0, *then E*_0_ *and E*_*B*_ *exist;*

*Case 4: if* ℱ_1_ *≤* 0 *and* ℱ_2_ *≤* 0, *then only E*_0_ *exists. The cases are depicted in Figure 2*.

**Proof**. We find an equilibrium solution (*T* ^*\**^, *C*^*\**^, *B*^*\**^, *A*^*\**^) of the system (1) from the steady state equations

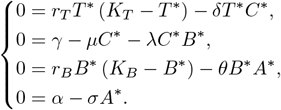

From the last equation we get that 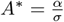, substituting it back to the third one we have

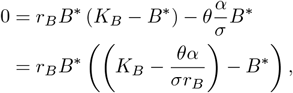

which leads to two possibilities for *B*^*\**^:

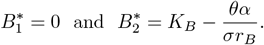

Since *B*(*t*) is nonnegative, 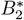 can only exist if 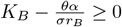, which is exactly the case when ℱ_1_ *≥* 0. From now on we suppose that ℱ_1_ > 0, i.e. 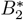 is positive. We will discuss the case of equality later. Now we substitute the possible *B* components of the equilibria into the equation for *C*:

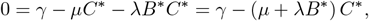

thus 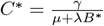, i.e.

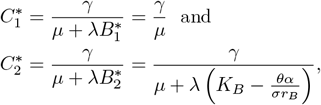

where for the existence of 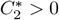 we also need ℱ_1_ *>* 0 to stand. Lastly, we substitute the derived *C*^*\**^ values into the first equation:

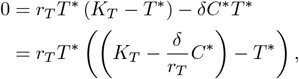

hence the possible *T* components of the equilibrium points are 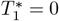 and 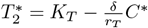, i.e.

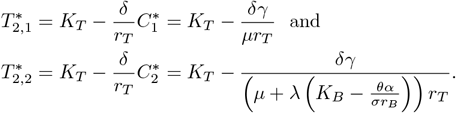

Since *T* (*t*) is nonnegative, 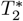 can only exist if 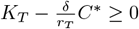, but similarly to 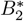, we also require the strict inequality, so 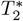 is positive. In the case of 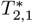, the condition 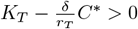 stands iff ℱ_2_ *>* 0, while in the case of 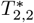, it is equivalent to ℱ_3_ *>* 0. For the existence of 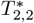, we also need ℱ_1_ *>* 0, inherited from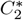.

We look separately at the cases when there is equality somewhere. If ℱ_1_ = 0, then 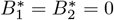, i.e. the bacteria are not present. If ℱ_2_ = 0 or ℱ_3_ = 0, then 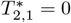 or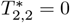, respectively, so in these situations the tumour is absent.

Hence, we obtain that if ℱ_1_ *≤* 0, then there is no positive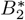. Furthermore, if ℱ_2_ *≤* 0 also, then a positive 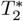 does not exist either, so the only possibly equilibrium is *E*_0_, otherwise *E*_0_ and *E*_*B*_ exist. If the condition ℱ_1_ *>* 0 stands, then there exist a positive 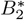. In this case, similarly, if ℱ_3_ *≤* 0, then there is no positive 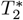, so only the equilibrium points *E*_0_ and *E*_*T*_ exist. Finally, if the condition ℱ_3_ *>* 0 holds, then there are two possible scenarios: if ℱ_2_*≤* 0, then there is no positive 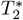 associated with *B* = 0, so only *E*_0_, *E*_*T*_ and *E*_coex_ exist, while in the other case, all four types of equilibria exist.

#### Proposition 2.4.

*There are just these five possible scenarios, since in Figure 2, the three lines* ℱ_1_ = 0, ℱ_2_ = 0 *and* ℱ_3_ = 0 *intersect in the same point*.

**Proof**. From ℱ_1_ = 0 we obtain that 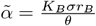, which substituted into ℱ_3_ = 0 gives us

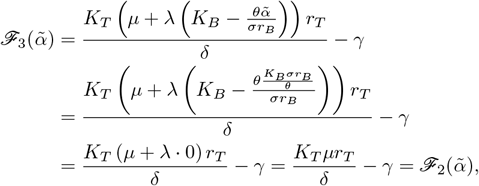

i.e. the lines ℱ_1_ = 0 and ℱ_3_ = 0 intersect on the line ℱ_2_ = 0.

So, we have seen that our system of equations (1) has four possible types of equilibria, and we have shown when each of them exists depending on the parameters of the equation system. In the following, we will determine the stability of these equilibria and prove the convergence of the solutions to the stable one. To achieve this, we use Theorem 1.3. from [20].

#### Theorem 2.5

([20]). *Suppose f is locally Lipschitz continuous with respect to x, uniformly in t. Let x*(*t*) *and y*(*t*) *be two differentiable functions such that*

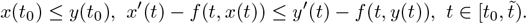

*Then we have x*(*t*) *≤ y*(*t*) *for every* 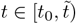. *Moreover, if x*(*t*) *< y*(*t*) *for some t, this remains true for all later times*.

Based on this theorem, we can prove the following comparison principle, a key tool in our proofs.

#### Proposition 2.6.

*Let f and g be locally Lipschitz continuous functions, and let x*(*t*) *and y*(*t*) *be two differentiable functions such that*

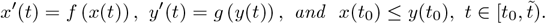

*Suppose that f* (*x*) *≤ g*(*x*) *holds for every x, then* 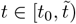.

**Proof**. Since *x*^*′*^(*t*) = *f* (*x*(*t*)) and *y*^*′*^(*t*) = *g* (*y*(*t*)), we see that

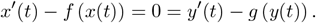

Since *f* (*x*) *≤ g*(*x*) holds for all *x*, it follows that

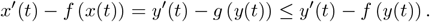

Then *f, x*(*t*) and *y*(*t*) satisfy the conditions of Theorem 2.5, from which we obtain *x*(*t*) *≤ y*(*t*) for all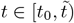,. Let ℝ^+^ and 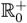 denote the set of the positive and nonnegative real numbers, respectively.

#### Theorem 2.7

(Convergence and stability). *In Case 1 the equilibrium E*_*coex*_, *in Case 2 the equilibrium E*_*T*_, *in Case 3 the equilibrium E*_*B*_, *and in Case 4 the equilibrium E*_0_ *is globally asymptotically stable in the sense that it is stable and attracts all solutions starting from*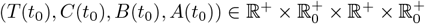. *Analogous results hold in the subspaces where either T* = 0 *or B* = 0, *according to Table 2*.

**Table 2:**
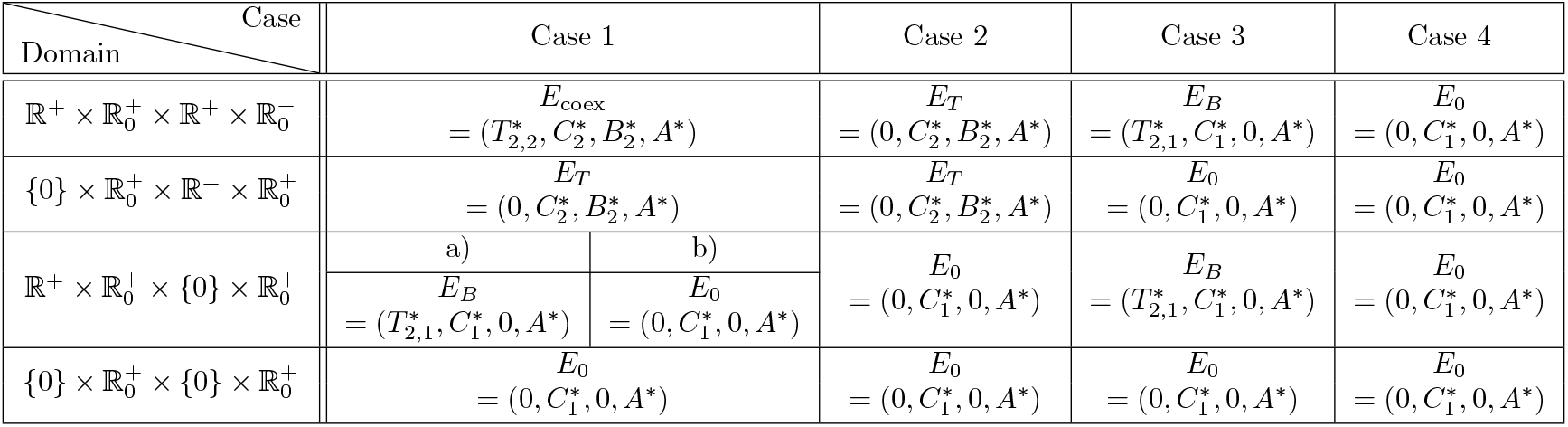
Possible stable equilibria of the equation system (1).

**Proof**. It is easy to see that for *T* (*t*_0_) = 0, the solution *T* (*t*) *≡*0, and for *B*(*t*_0_) = 0, the solution *B*(*t*) *≡*0. This implies that these two functions will always take the value 0, no matter how the parameters of the system of equations are chosen, so they cannot converge to positive equilibrium situations. If *B*(*t*_0_) = 0, we also have to take into consideration the sign of ℱ_2_ in Cases 1 and 2. Because of *B*(*t*) *≡* 0, the positive equilibrium 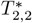 cannot exist, but for the existence of a positive 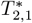 we need the condition ℱ_2_ *>* 0, which we do not have here. In Case 1, we already did a separation (Case 1 a) and b)), and in Case 2, ℱ_1_ *>* 0 and ℱ_3_ *≤* 0 imply ℱ_2_ *≤* 0, so only 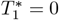exists.

Since in the case of *T* (*t*_0_) = 0 or *B*(*t*_0_) = 0 the phenomenon under consideration would not occur (i.e. we would not have bacteria sabotaging the cancer treatment), we can assume that *T* (*t*_0_), *B*(*t*_0_) *∈* ℝ^+^ and 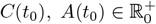, i.e. we are working on the domain 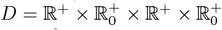, which is the most interesting case. The other scenarios listed in Table 2 can also be proved.

Consider first the equation *A*^*′*^(*t*) = *α − σA*(*t*) for the function *A*(*t*), which is an inhomogeneous linear equation, so it has the solution

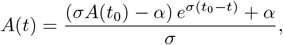

from which we get

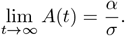

From that we can see the *A* component of the equilibrium point: 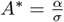, and that *A*(*t*) converges to *A*^*\**^, e.g. 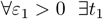:

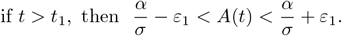

Next we look at the equation *B*^*′*^(*t*) = *r*_*B*_*B*(*t*) (*K*_*B*_ *− B*(*t*)) *− θB*(*t*)*A*(*t*) which describes the size of the bacteria population. Let

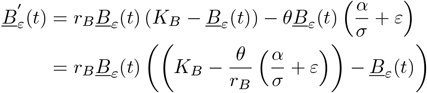

and

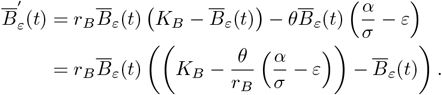

The equations for *B*_*ε*_(*t*) and 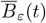 will still be logistic if 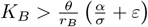, so the solutions are:

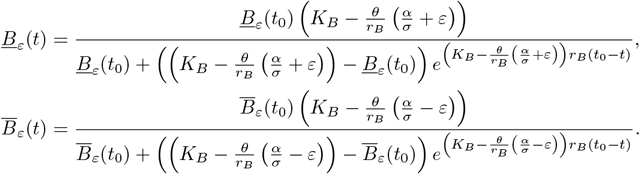

Since for *t > t*_1_ the inequalities 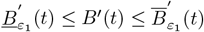 stand, we get 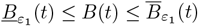 from Proposition 2.6 if we set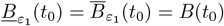. Note that

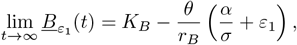

And

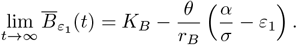

This implies that

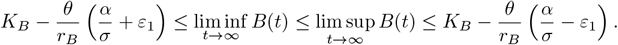

Since these inequalities hold for all *ε*_1_ *>* 0, we get that if 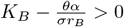e.g. ℱ_1_ *>* 0 stands, then

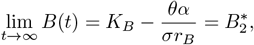

Otherwise

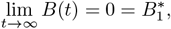

so *B*(*t*) converges to one of the *B* components of the equilibria characterized in Table 2.

Next we consider the equation describing the change of the concentration of the chemotherapy drug as follows: *C*^*′*^(*t*) = *γ − µC*(*t*) *− λC*(*t*)*B*(*t*). There are two cases based on which limit point of *B*(*t*) we use.

1. If 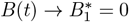, e.g. 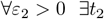:

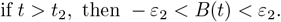

Let

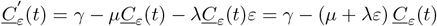

and

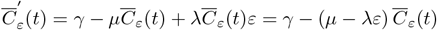

The equations for 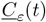 and 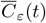 will still be inhomogeneous linear equations if *µ − λε >* 0, so their solutions are:

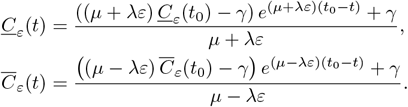

Since the inequalities 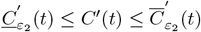 stand when *t > t*_2_, we get 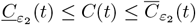 from Proposition 2.6 if we set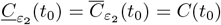. Note that

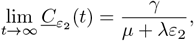

and

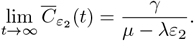

Since *ε*_2_ *>* 0 was arbitrary, thus

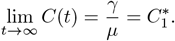

2. If 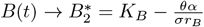, e.g. 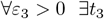:

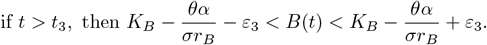

Similarly to the previous case, now we use these boundaries and the functions 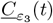 and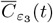. Then applying Proposition 2.6. again, we get that in this case

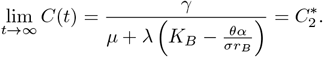

So we can see that *C*(*t*) also converges to one of *C* components of the equilibria in Table 2.

Last we consider the equation *T* ^*′*^(*t*) = *r*_*T*_ *T* (*t*) (*K*_*T*_ *− T* (*t*)) *− δT* (*t*)*C*(*t*) describing the change of the tumour size. In a completely analogous way to the calculations done earlier for *B*(*t*), we obtain the following.

1. In the case of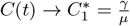, if 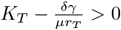e.g. ℱ_2_ *>* 0, then

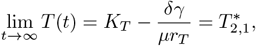

Otherwise

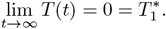

2. In the case of 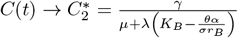, if ℱ_3_ *>* 0, then

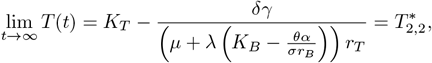

Otherwise

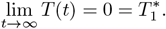

So *T* (*t*) also converges to one of the *T* components of the equilibria in Table 2.

Thus, we obtain that all four functions converge to the corresponding components of an equilibrium characterized in Table 2., i.e. the solutions converge. In the following, we investigate the global asymptotic stability of the equilibrium situations.

The stability of an equilibrium point is shown using *ε − δ* neighborhoods. Since we have a final dimensional system, we can show that componentwise, and for that, we will use the pinching functions again. Here we investigate the stability of 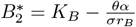, the rest can be proved in a similar way. Let *ε >* 0 be given and let 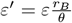, then

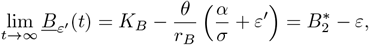

and

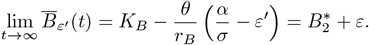

Let 0 *< δ < ε* be arbitrary and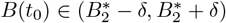. Now consider the functions 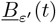 and 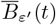starting from the points 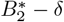and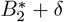, respectively, i.e.

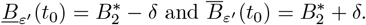

We know that the positive equilibrium of a logistic equation is asymptotically stable, moreover, the solution converges to it monotonically. This means that the 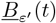 function starting from the point 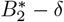 will converge to 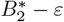 in a monotonically decreasing way, and the function 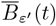starting from the point 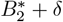 will converge to 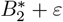 in a monotonically increasing way. Thus both of these functions stay in the interval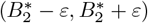. Since the conditions of Proposition 2.6 hold, we get that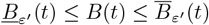, so *B*(*t*) also stays in the interval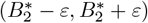, which we wanted.

In the remaining cases, the stability of a positive component of an equilibrium can be derived from analogous calculations. If we consider a 0 component, e.g. 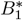, then we use 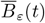 and the constant 0 function as the pinching functions. Thus the previously characterized equilibria of the system are all globally asymptotically stable.

#### Remark 2.8

(Bifurcations). A transcritical bifurcation is the situation when, as a parameter varies, a stable and an unstable equilibrium come together and exchange stability at the critical point (in higher dimensional systems, this happens in some subspace), i.e., there is a stable and an unstable equilibrium both before and after the bifurcation, but their role is reversed ([21]).

In our model, transcritical bifurcations are observed at the points where the sign of a threshold parameters changes. Typically, the positive component of an equilibrium becomes zero at the critical point and changes sign, but then it moves outside of the biologically meaningful domain, so we only track it until reaching 0. Since we can explicitly describe the equilibria and their stability, we can also identify these bifurcations: both the meeting point of two equilibria, as well as how their stability changes. Overall we find the following bifurcations.

- Between Case 1 a) and Case 1 b), i.e. when the sign of ℱ_2_ changes. If we increase *γ*, then the positive 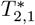 will decrease, and eventually become 0. So when ℱ_2_ becomes nonpositive, then *E*_*B*_ merges into *E*_0_.
- Between Case 1 b) and Case 2, i.e. when the sign of ℱ_3_ changes. If we increase *γ*, then the positive 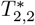 will decrease, and eventually become 0. So when ℱ_3_ becomes nonpositive, then *E*_coex_ merges into *E*_*T*_. The same happens if we increase *α* instead of *γ*.
- Between Case 3 and Case 4, i.e. when the sign of ℱ_2_ changes. If we increase *γ*, then the positive 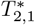 will decrease, and eventually become 0. So when ℱ_2_ becomes nonpositive, then *E*_*B*_ merges into *E*_0_.
- Between Case 1 a) and Case 3, i.e. when the sign of ℱ_1_ changes. If we increase *α*, then the positive 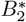 will decrease, and eventually become 0. In this case, two transcritical bifurcations happen simultaneously: when ℱ_1_ becomes nonpositive, then *E*_coex_ and *E*_*T*_ merge into *E*_*B*_ and *E*_0_, respectively.
- Between Case 2 and Case 4, i.e. when the sign oℱ_1_ changes. If we increase *α*, then the positive 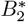 will decrease, and eventually become 0. So when ℱ_1_ becomes nonpositive, then *E*_*T*_ merges into *E*_0_.

## 3 Simulations and data fitting

### 3.1 Simulations

We derived that the treatment has four possible outcomes characterized by the signs of the threshold parameters. These outcomes are illustrated in Figures 4.-7., which were made with the following parameter values: *r*_*T*_ = 0.4, *K*_*T*_ = 2.62, *δ* = 1.15, *µ* = 2.6, *λ* = 2, *r*_*B*_ = 1.81, *K*_*B*_ = 1.81, *θ* = 1.085, *σ* = 0.7, and we have *α ∈* {1.39, 3.06} and *γ ∈* {1.24, 3.85}. In the colored squares on the left side of the figures (*α ∈* [0, 6], *γ ∈* [0, 6]) the red points denote the current (*α, γ*) values fow which the simulation was run.

### 3.2 Data fitting

In [17], colon cancer cells were implanted under the skin of mice and their growth was monitored, and we use this to test a specified version of our model. Tumour growth was measured over a nine-days interval, while the tumour was treated with Gemcitabine on the fourth and ninth days, as described in the figure caption, and the relative tumour volume (proportional to the first day volume) was plotted as a function of time (Figure 1.B in the article). However, their image description and the measured data is not completely in sync, hence we will fit the timing of the second dose of chemotherapy to the observed data. In this experiment no additional antibiotic treatment was given, hence we omit the *A*-equation, and consider

**Fig. 1:**
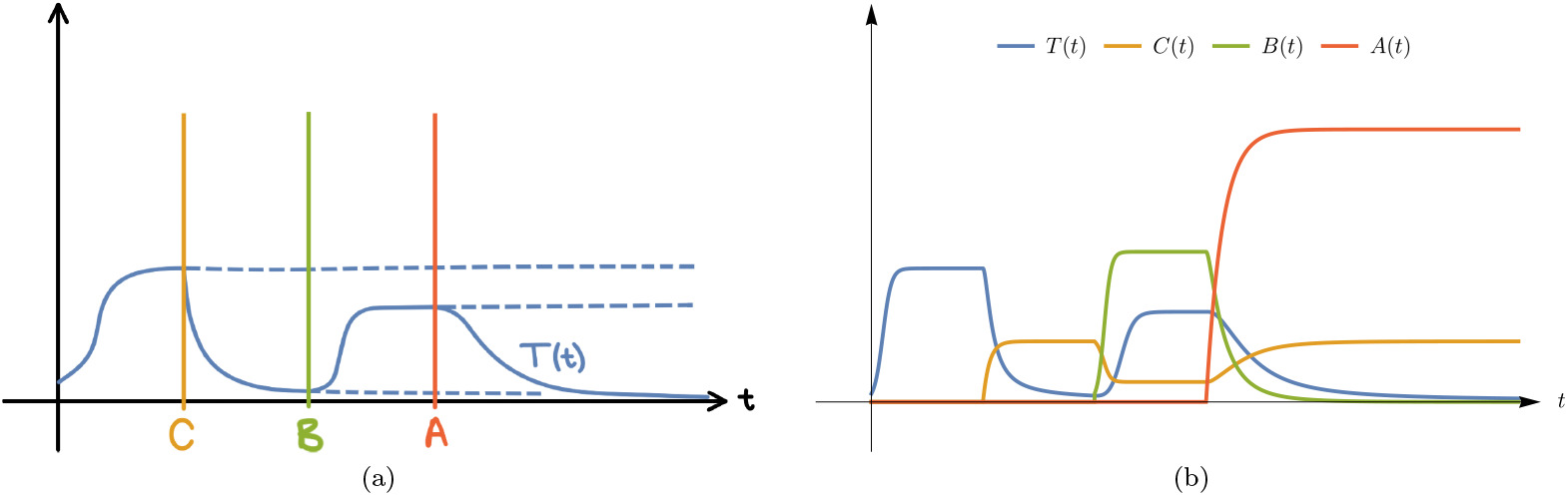
(a) Based on biological intuition, a possible scenario for the change of tumour size (*T* (*t*)) in response to different externalities. Here, at time *C* the chemotherapy treatment is started, decreasing the tumour size. At time *B* the patient is infected by the bacteria, and at time *A* the patient receives the antibiotics treatment. (b) Recreation of the same scenario by a numerical simulation using our mathematical model. Additional to variation of the tumour size, we plot the drug concentrations and bacterial density as well.

**Fig. 2:**
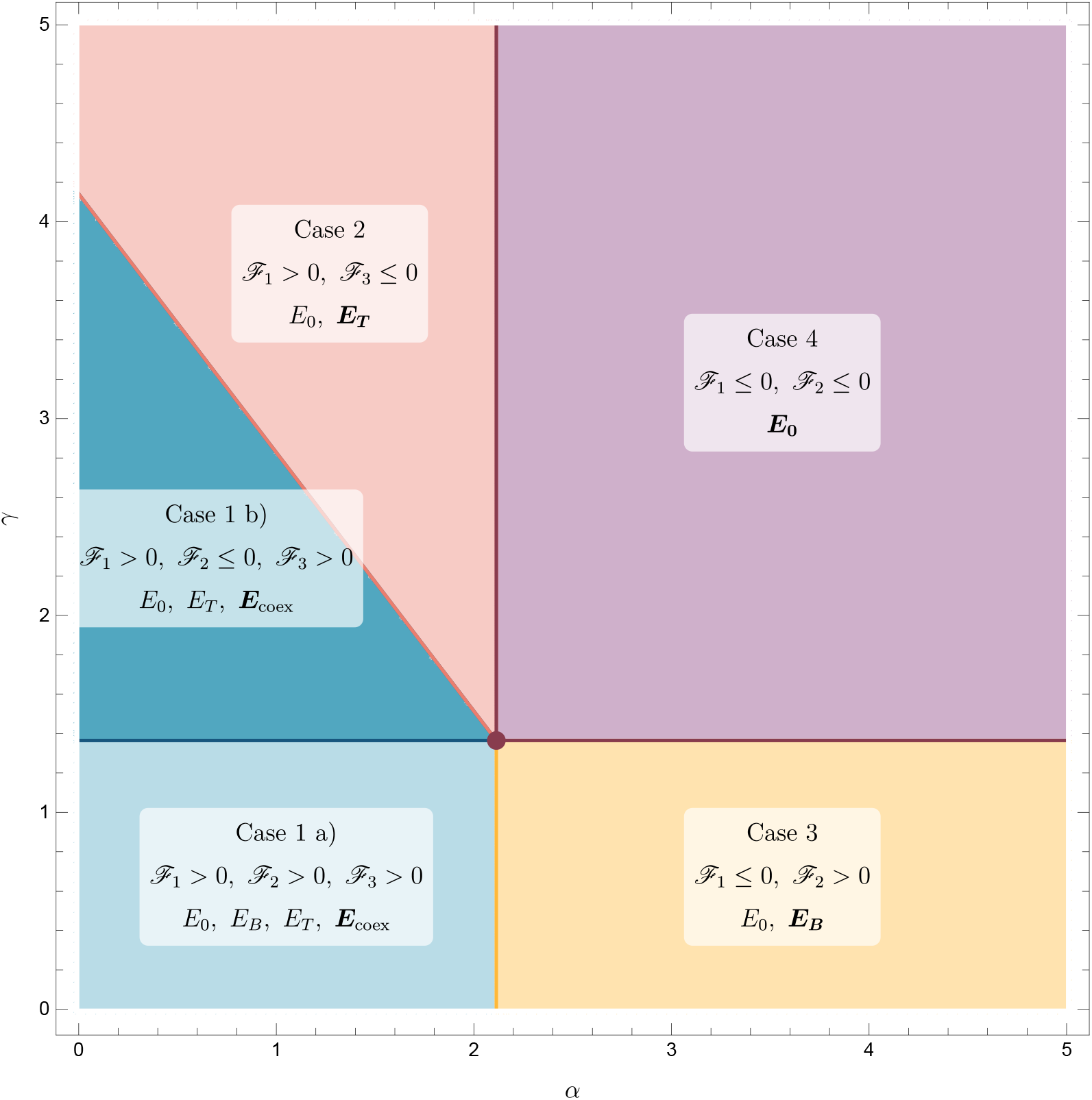
Illustration of the five possible cases in Theorem 2.3, in the parameter plane of the two drug doses. The equilibria with bold are the stable ones discussed in Theorem 2.7. The equations of the (half) lines on the boundaries: vertical: ℱ_1_ = 0, horizontal: ℱ_2_ = 0, tilted on the left: ℱ_3_ = 0. The figure was made using the parameters *r*_*T*_ = 1.71, *K*_*T*_ = 1.61, *δ* = 3.59, *µ* = 1.78, *λ* = 2, *r*_*B*_ = 1.81, *K*_*B*_ = 1.81, *θ* = 1.085, *σ* = 0.7 and *α, γ ∈* [0, 5].

**Fig. 3:**
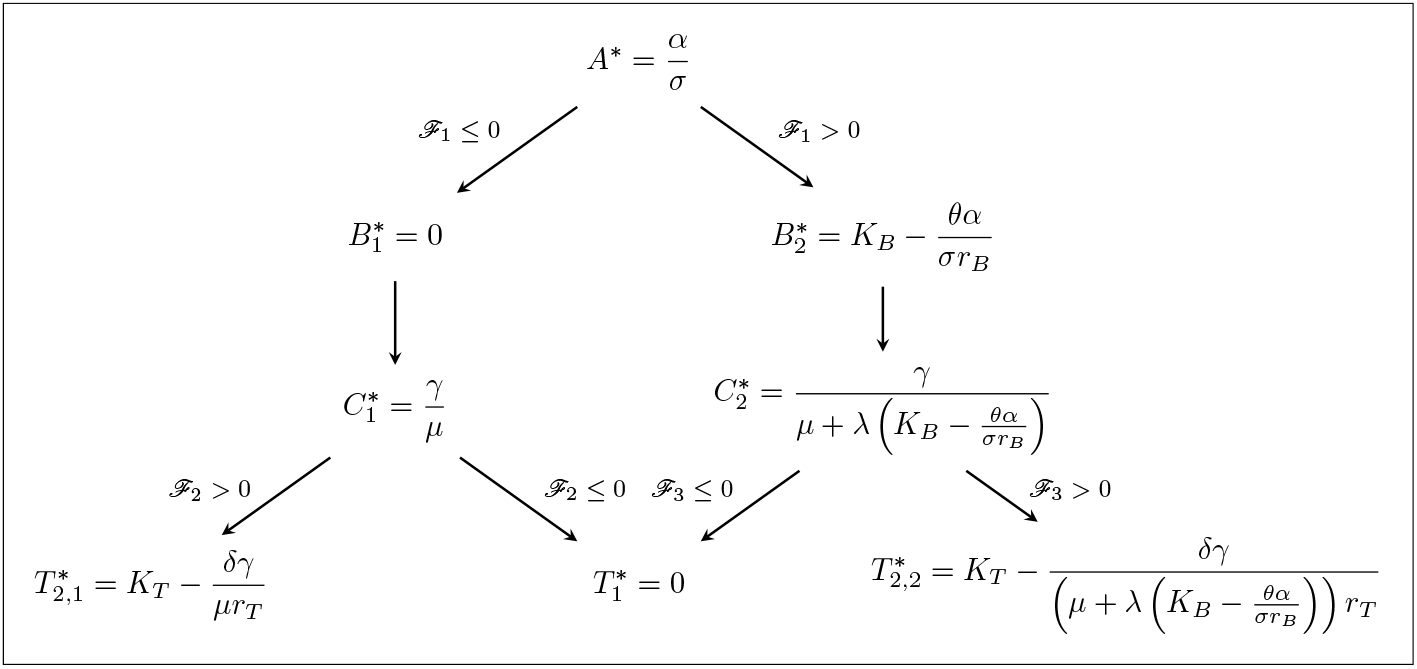
Derivation of the possible coordinates of the equilibria depending on the signs of the threshold parameters.

**Fig. 4:**
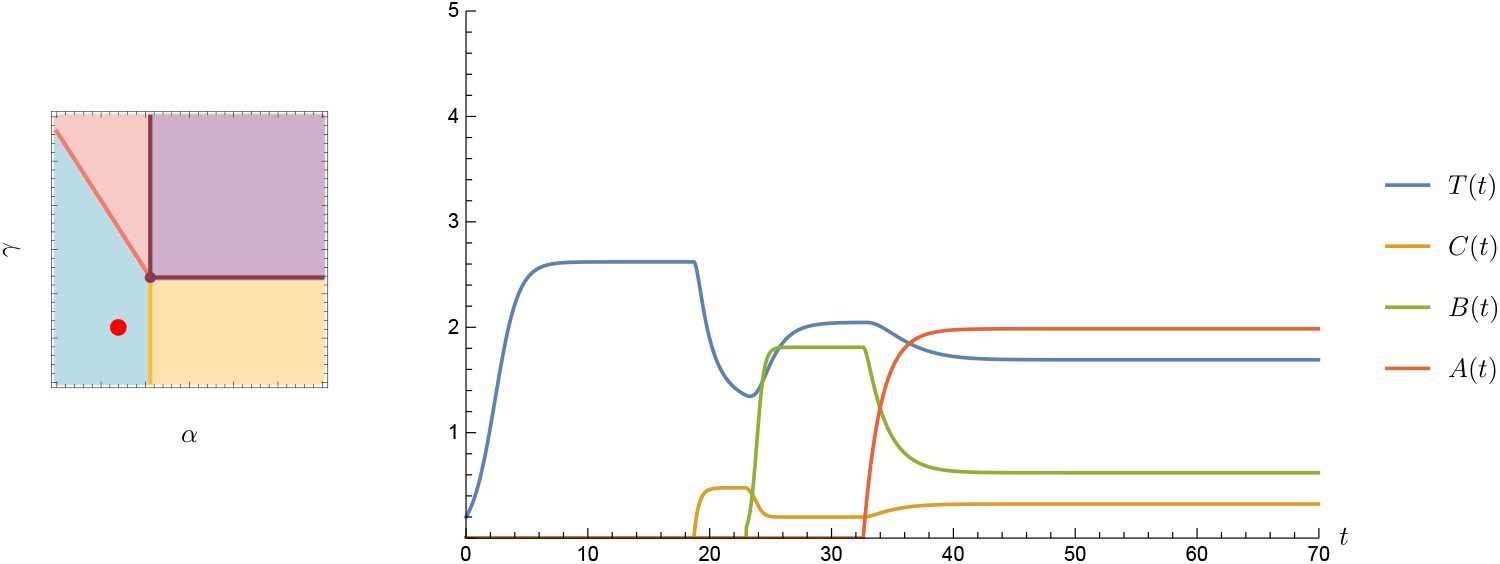
Both the bacteria population and the tumour survive. Here *α* = 1.39 and *γ* = 1.24.

**Fig. 5:**
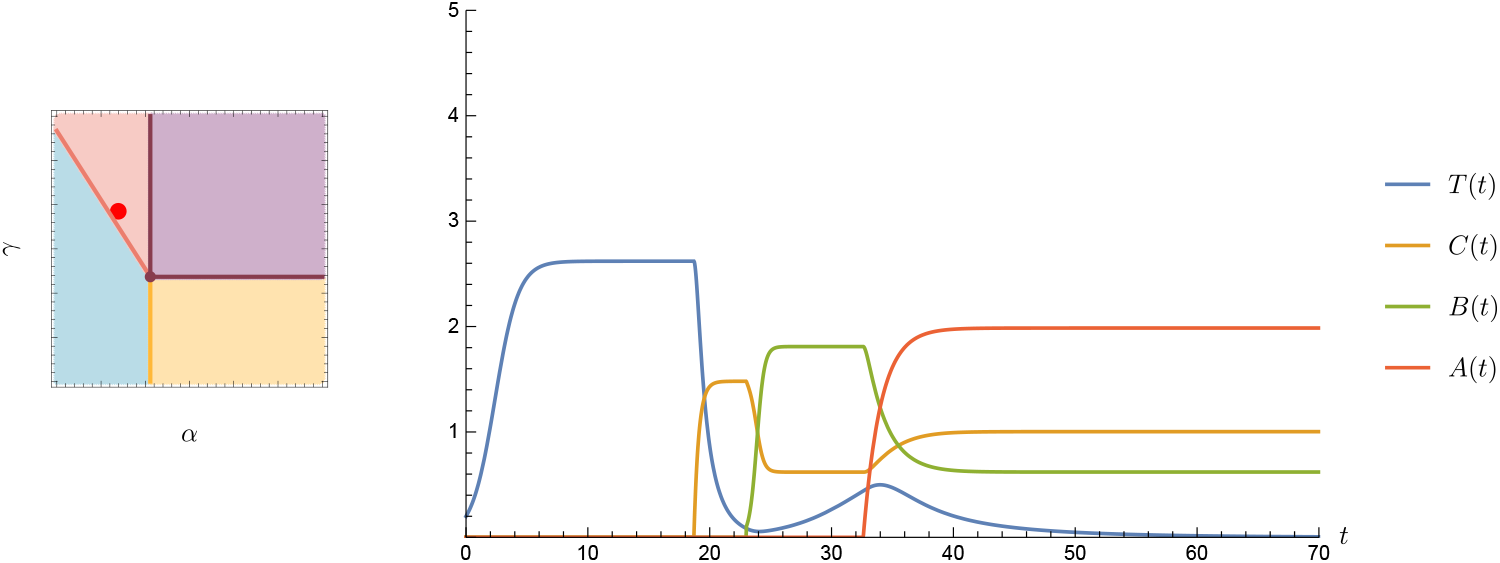
The bacteria survive, but the tumour dies out. Here *α* = 1.39 and *γ* = 3.85.

**Fig. 6:**
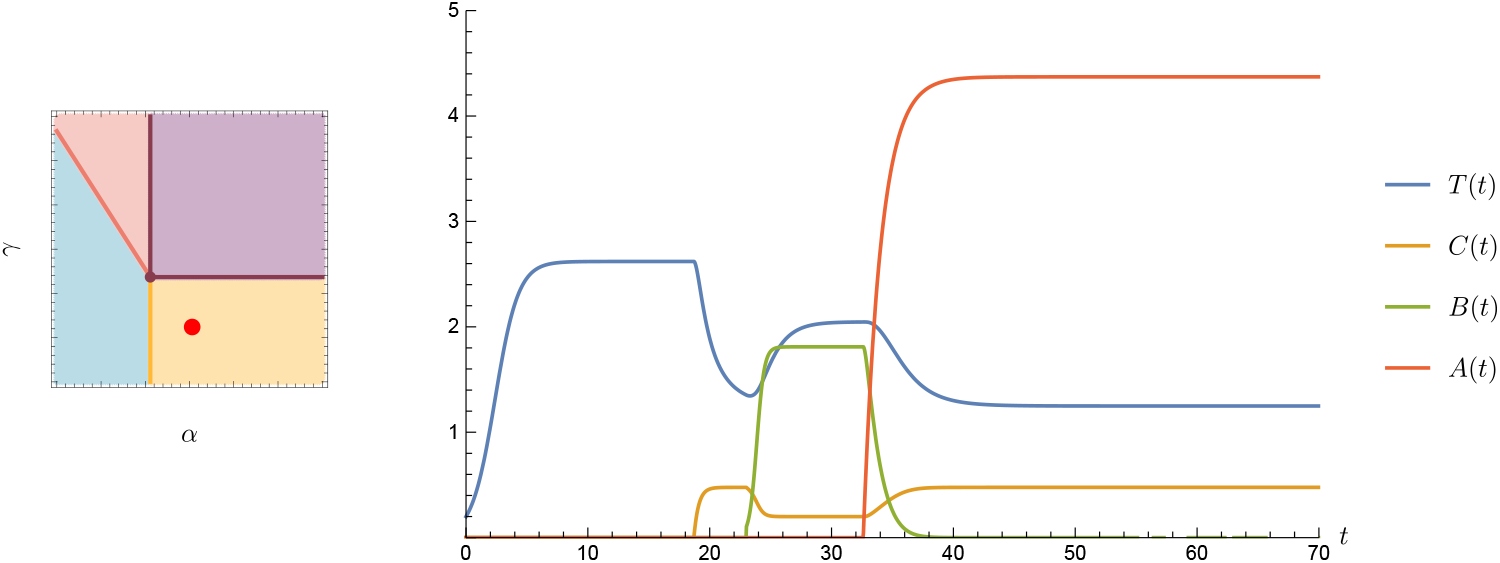
The tumour survives, but the bacteria die out. Here *α* = 3.06 and *γ* = 1.24.

**Fig. 7:**
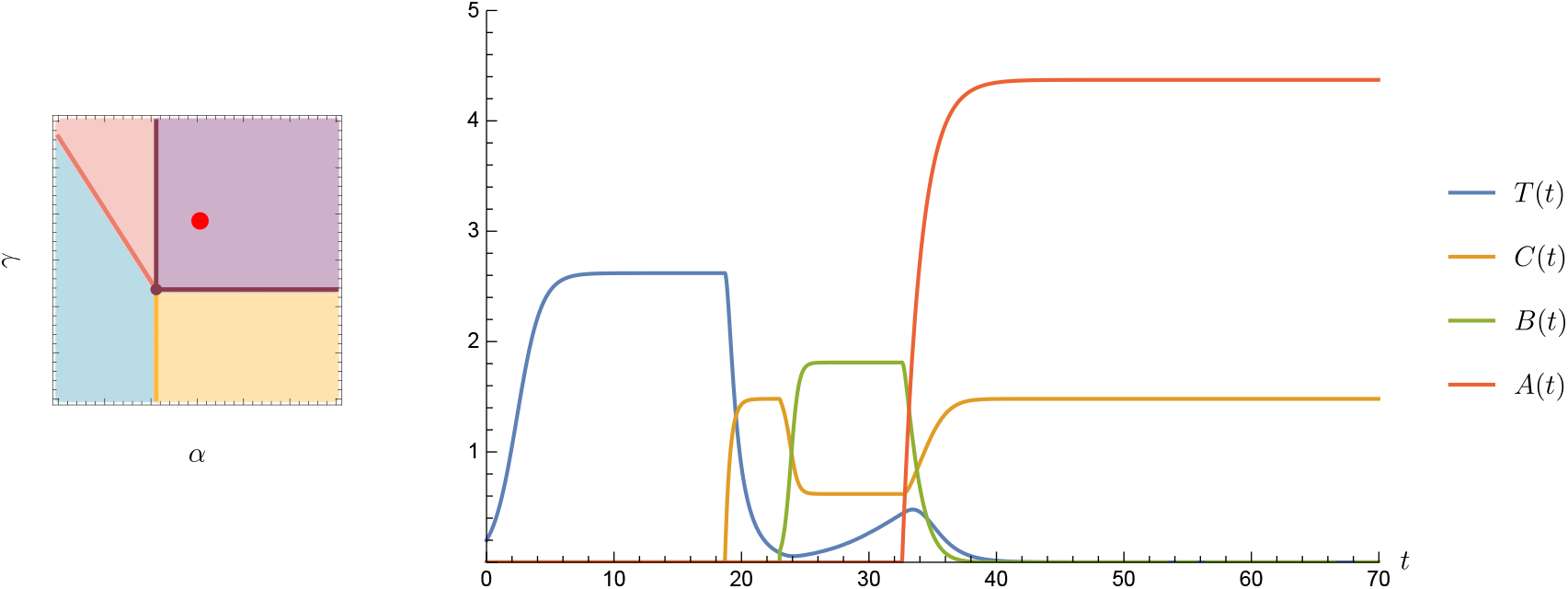
Both the tumour and the bacteria population die out. Here *α* = 3.06 and *γ* = 3.85.

**Fig. 8:**
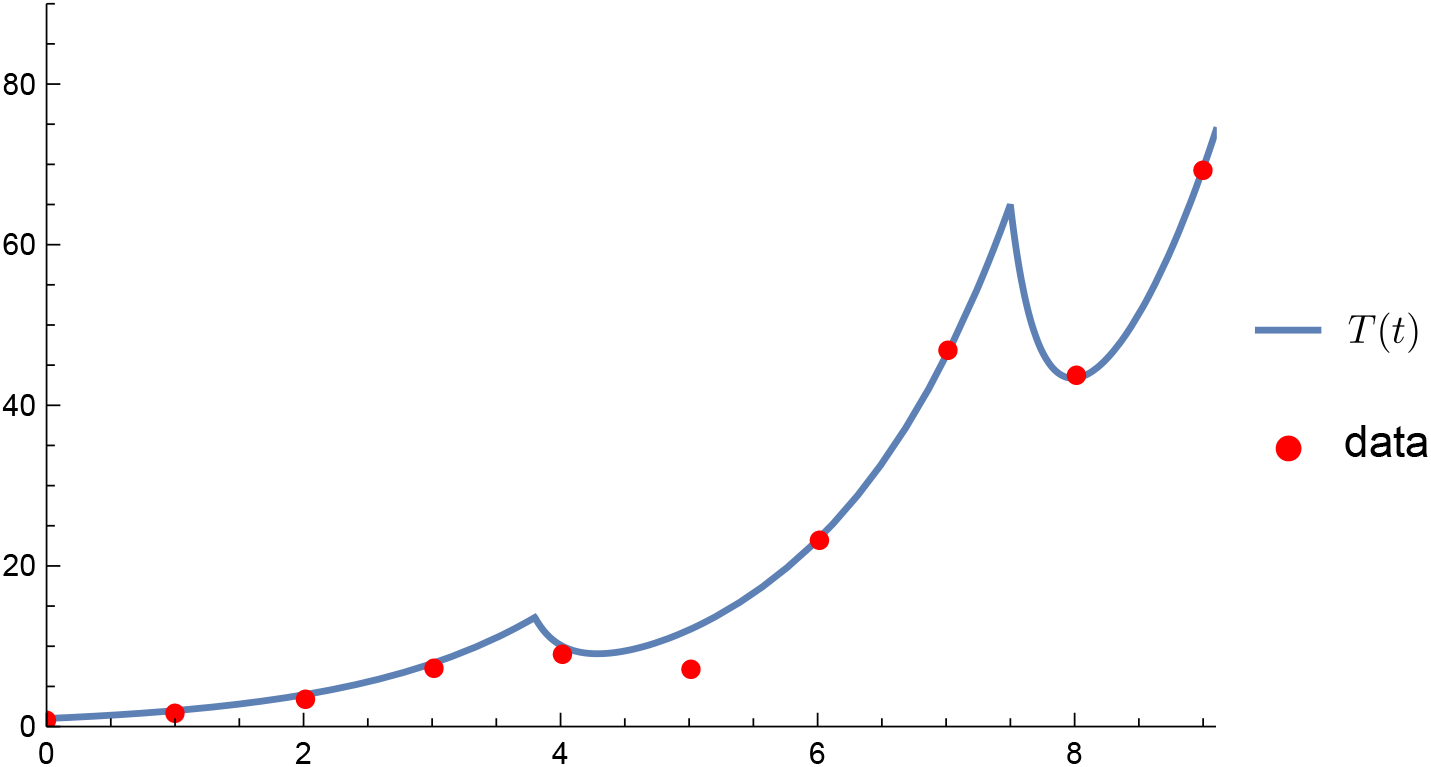
The best fit of the function *T* (*t*) to the measured data. The optimized parameter values are: *r* = 0.6867, *K*_*T*_ = 51831.9, *δ* = 0.06312, *φ* = 0.1325, and the precise times of the chemotherapeutic treatment administration are *t*_1_ = 3.8, *t*_2_ = 7.5.

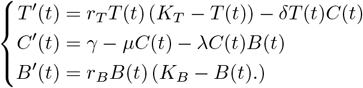

For convenience, we rewrite the first equation as

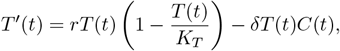

where *r* = *r*_*T*_ *K*_*T*_. During the experiment, which was a relatively short time period, the bacteria population size is assumed to be constant *B*_0_. As in the experiment, the chemotherapy drug will not be administered continuously, but will be given at times *t*_1_ and *t*_2_. Thus, except for these two time points, the change of the chemotherapy drug will described with the equation

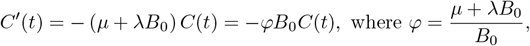

and then we can think of *φ* as a kind of combined rate of decay. Having these modifications, we consider our new system of impulsive differential equations:

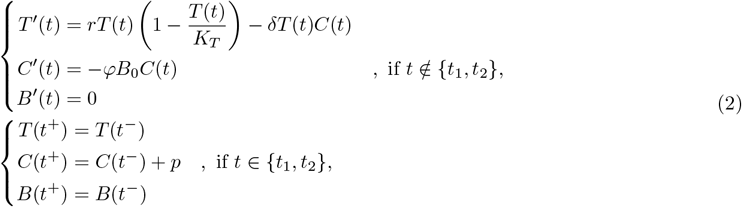

where 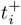 is the time at which the *p* dose of chemotherapy drug is administered, and 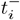 is the last moment immediately before administration for *i* = 1, 2.

The parameters of the system of equations (2) were optimized using the FindFit procedure of Wolfram Mathematica software (version 12.2), based on the method of least squares. Since it is not known at which time of day the tumour volume was measured or the chemotherapy drug was administered on a given day, the time of drug administration is also a parameter to be optimized. Thus, *r, K*_*T*_, *δ, φ, t*_1_, *t*_2_ are the parameters to be estimated, and the following constraints were imposed in order to obtain a good and feasible fit:

- all parameters are positive,
- the constant amount of the bacteria is *B*_0_ = 22 units (this can be scaled by *φ*, hence this value is actually irrelevant),
- to match the possible dosage times in the experiment, we restrict 3.8 *< t*_1_ *<* 4.5 and 6.8 *< t*_2_ *<* 7.5.
- for the tumour, the capacity is *K*_*T*_ *>* 400 units,
- the dose of the chemotherapy drug is *p* = 45 units (its effect is scaled by *δ* hence the exact value is irrelevant).

In the best fit, the least squared error was 24.7869. The data series and the graph of the best fit are shown in Picture 8.

## 4 Generalizations and robustness

### 4.1 More universal growth rate

When setting up the equation system (1), we made several assumptions, one of which was that the growth of the tumour and the bacteria population was modelled by a logistic equation. Instead, however, we can use more general functions with some simple – and biologically realistic – assumptions that ensure that our previous results remain true.

Let *f* (*T* (*t*)) be the growth rate at time *t* for the tumour size *T* (*t*), then our equation describing the tumour is

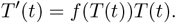

We need *f* to be a continuous, monotonically decreasing function, but this is a natural assumption, since the larger the tumour, the less nutrients and resources a tumour cell has. Also, we assume that *f* has a positive root, *T* ^0^. Then *f* (*T* ^0^) will be the nontrivial equilibrium state of the equation, since *T* ^*′*^(*t*) = *f* (*T* ^0^)*T* (*t*) = 0. For *T* (*t*) *< T* ^0^, *f* (*T* (*t*)) is positive, so the tumour increases, while for *T* (*t*) *> T* ^0^, *f* (*T* (*t*)) is negative, so the tumour shrinks, thus *f* (*T* (*t*)) indeed tends to *f* (*T* ^0^). For the bacteria, a similar condition can be used to define a general growth rate *g*(*B*(*t*)). Then our system of equations is modified to the following form:

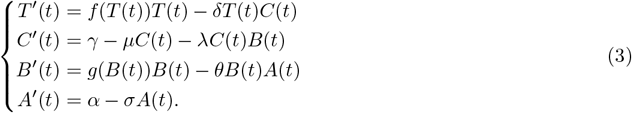

As in the case of the equation system (1), it can be seen that for nonnegative initial values, the solutions of the equation system (3) will also be nonnegative and bounded. The statements of Theorem 2.3. and Theorem 2.7. hold, i.e., in the case of more general functions *f* and *g*, equilibrium points exist and the solutions always converge to one of these. The proofs can be carried out in almost the same way in both cases. In the proof of Theorem 2.3. we exploit the fact that *f* and *g* are monotonically decreasing and both of them have a positive root. Consider, for example, the step when we substitute back the *C* component of the equilibrium (*C*^*\**^) into our equation for the tumour:

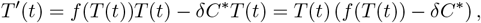

then again there can be at most one positive equilibrium, since *f* (*T* (*t*)) *− δC*^*\**^ also has one zero due to the conditions on *f*. To prove convergence, as in the proof of Theorem 2.7., one can use the previous comparison principle (we can squeeze the function between *±ε* versions) and Proposition 2.6 due to the continuity of *f* and *g*.

Examples for possible *f* (and similarly for *g*) functions:

- logistic equation: *f* (*T* (*t*)) = *r*_*T*_ (*K*_*T*_ *− T* (*t*)),
- Richards’s growth model, which is a generalization of the logistic one: 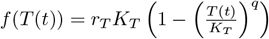where *q >* 0.
- Gompertz-function: an other commonly used growth model, where the growth of the tumour is described by the function 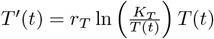,from which we get 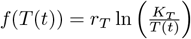.

### 4.2 Pharmacokinetics

During setting up the equation system (1), we assumed that the antibiotic drug kills bacteria in direct proportion to its concentration. In reality, however, this is not necessarily a linear relationship, and the dose-response curves often given by Hill functions:

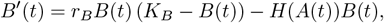

where 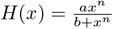 is a Hill-function. Similarly, we can replace the *−δT* (*t*)*C*(*t*) term for cytotoxic effect by the Hill-type term

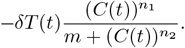

In this case, however, using a mass action term gave a better fit to data for the experiment described in Section 3.2.

### 4.3 Bacteria population dynamics

In our model we implicitly assumed that although the bacteria metabolize the chemotherapy drug, it does not substantially influence the growth rate of the bacteria population. We found no evidence for this in [17], but we can not exclude this possibility. In this case, an additional term appears in the equation describing the change in the enhanced growth rate of the bacteria population:

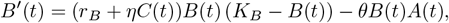

where it is assumed that the bacteria benefit in proportion to the amount of chemotherapeutic drug at a given time, and *η >* 0 is the parameter describing the extent of this benefit.

### 4.4 Impulsive drug doses

As we have done in Section 3.2., instead of giving a continuous dose of medicines, we can prescribe periodically taken medication. Then we write the *γ*(*t*) and *α*(*t*) functions in the following forms:

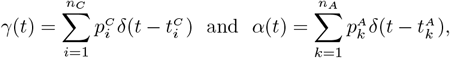

where *δ*(*t*) is the Dirac-delta, 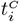 and 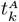 are the times when the patient takes the chemotherapy drug and the antibiotics, respectively, and 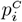 and 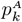 are the given doses corresponding to those times. This approach requires further consideration, for example, optimizing the time intervals between subsequent medications. By numerical simulations, we can confirm that similarly to the continuous dosage case, the four possible outcomes described earlier are achievable. The similar dynamical behaviour of the continuous and impulsive treatment models is highlighted by two examples shown in Figure 9.

**Fig. 9:**
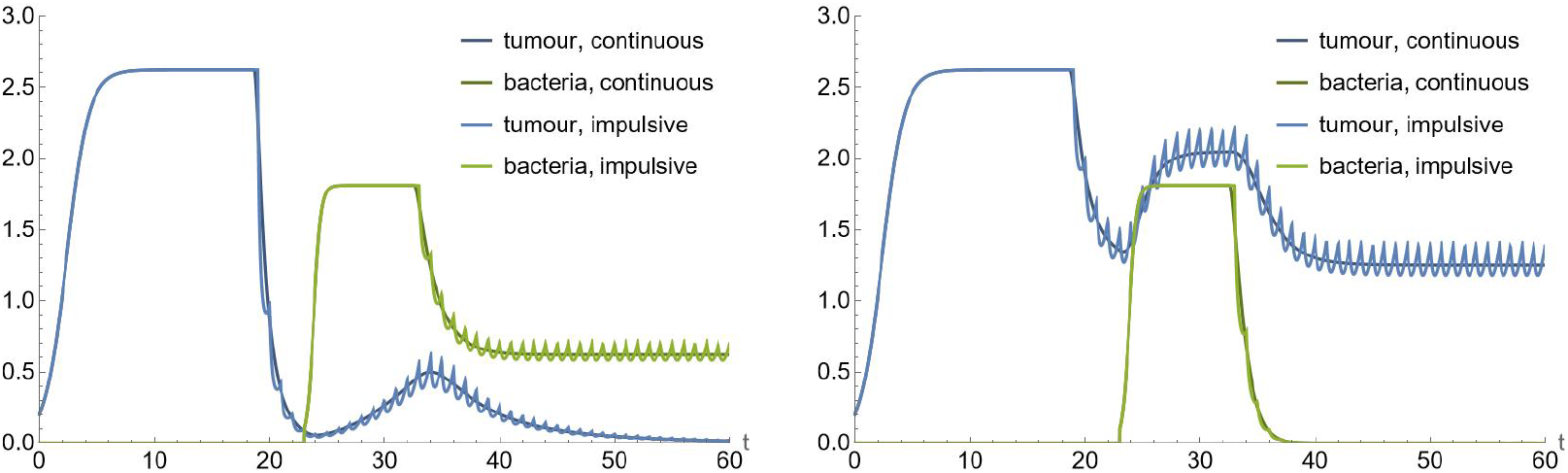
Comparison of treatment outcomes using continuous and impulsive drug dosage. In the impulsive case, the same amount of drug is given in each time unit, but all at once at integer time points. The left figure depicts a situation when the bacteria population persists, but the tumour is eradicated, while in the right figure the tumour remains despite the bacteria die out. These scenarios correspond to Case 2 and Case 3, respectively, and the plots show the same dynamics for the continuous and the impulsive treatment.

## 5 Conclusions

Our work was motivated by a recent biological discovery that certain bacteria can metabolize anti-cancer chemotherapy drugs. Therefore, if the bacteria are present in a treated patient, they have the potential to sabotage the treatment, as the metabolized drug cannot exert its anti-tumour effect. In the presence of sabotaging bacteria, it may be recommended to supplement the treatment with antibiotics. To describe this phenomenon, we constructed a mathematical model consisting of four differential equations was constructed, following the variation of the size of the tumour, the concentration of the chemotherapeutic drug, the size of the bacteria population and the concentration of the antibiotic drug. The model expresses the interactions between these components.

We introduced three compound threshold parameters and shown that their signs characterize the existence and stability of the equilibria. We found four possible outcomes, which we explicitly described: both the tumour and the bacterial population die, the tumour dies but the bacteria survive, the tumour survives even though the bacterial population dies, or neither the tumour nor the bacterial population dies. These outcomes are guaranteed by the global asymptotic stability of the corresponding equilibrium. The proof is based on the iterative application of a comparison principle. We highlighted these possibilities in a bifurcation diagram, and described the types of bifurcations in the transitions between the different cases, as single and simultaneous transcritical bifurcations.

Our model was able to reproduce a biological experiment, where rectal cancer tumour was grown in mice and then treated with the chemotherapeutic drug Gemcitabine. The experiment compared the tumour growth of bacteria in mice where the bacteria that metabolize the drug were present or absent. Since the drug was administered at specific time points, the model equations were augmented with corresponding impulsive terms. Parameter estimation was performed using data extracted from [17] and a relatively good fit was obtained. Our model fitting was able to estimate the precise timing of the treatments that was not reported in the biological publication.

We investigated the robustness of our findings by considering possible generalizations of the model and showed how the results extend to such situations. Our model has limitations: we assume homogeneous spatial distribution and a well-mixed system of tumour cells, bacteria and drugs. Potential future research can incorporate this and other important biological factors. In particular, a promising future research direction is the application of a control theory approach to optimize the combined treatment of chemotherapy and antibiotics when the saboteur bacteria are present. The model we presented here contributes to the better understanding of the role of intratumour bacteria in the success or failure of cancer treatment.

## CRediT authorship contribution statement

Anna Geretovszky: Conceptualization, Formal analysis, Methodology, Visualization, Writing - original draft, Writing - review & editing.

Gergely Röst: Conceptualization, Formal analysis, Funding acquisition, Methodology, Supervision, Writing - review & editing.

## Declaration of Competing Interest

The authors declare that they have no known competing financial interests or personal relationships that could have appeared to influence the work reported in this paper.

## Acknowledgement

AG was supported by the ÚNKP-23-2 - New National Excellence Program of the Ministry for Culture and Innovation from the source of the National Research, Development and Innovation Fund (grant number ÚNKP-23-2-SZTE-586), and TKP-2021-EGA-05, 2022-2.1.1-NL-2022-00005. GR was supported by RRF-2.3.1-21-2022-00006 and KKP 129877.

## Notes

### Competing Interest Statement

The authors have declared no competing interest.

